# Muscle Coordination Matters: Insights into Motor Planning using Corticospinal Responses during Functional Reaching

**DOI:** 10.1101/2023.05.15.540531

**Authors:** Thomas E Augenstein, Seonga Oh, Trevor A Norris, Joshua Mekler, Amit Sethi, Chandramouli Krishnan

**Affiliations:** NeuRRo Lab, Department of Physical Medicine and Rehabilitation, Michigan Medicine, Ann Arbor, MI, 48108 USA; Department of Robotics, University of Michigan, Ann Arbor, MI, 48109 USA; Physical Medicine and Rehabilitation, Michigan Medicine, Ann Arbor, MI, 48109 USA; Neurology, Michigan Medicine, Ann Arbor, MI, 48109 USA; Department of Occupation Therapy, University of Pittsburgh, Pittsburgh, PA, 15219 USA

## Abstract

The central nervous system (CNS) moves the human body by forming a plan in the primary motor cortex and then executing this plan by activating the relevant muscles. It is possible to study motor planning by using noninvasive brain stimulation techniques to stimulate the motor cortex prior to a movement and examine the evoked responses. Studying the motor planning process can reveal useful information about the CNS, but previous studies have generally been limited to single degree of freedom movements (*e.g.,* wrist flexion). It is currently unclear if findings in these studies generalize to multi-joint movements, which may be influenced by kinematic redundancy and muscle synergies. Here, our objective was to characterize motor planning in the cortex prior to a functional reach involving the upper extremity. We asked participants to reach for a cup placed in front of them when presented with a visual “Go Cue”.

Following the go cue, but prior to movement onset, we used transcranial magnetic stimulation (TMS) to stimulate the motor cortex and measured the changes in the magnitudes of evoked responses in several upper extremity muscles (MEPs). We varied each participant’s initial arm posture to examine the effect of muscle coordination on MEPs. Additionally, we varied the timing of the stimulation between the go cue and movement onset to examine the time course of changes in the MEPs. We found that the MEPs in all proximal (shoulder and elbow) muscles increased as the stimulation was delivered closer to movement onset, regardless of arm posture, but MEPs in the distal (wrist and finger) muscles were not facilitated or even inhibited. We also found that facilitation varied with arm posture in a manner that reflected the coordination of the subsequent reach. We believe that these findings provide useful insight into the way the CNS plans motor skills.

## Introduction

An essential part of daily life for many humans is manipulating their physical environment in reaction to sensory stimuli. For instance, someone who feels thirsty will see a glass of water and reach forward to pick it up and drink it. At a neuromuscular level, this process starts with sensory (*e.g.,* visual) stimuli producing afferent neural signals that the central nervous system (CNS) uses as a cue to initiate the movement. A motor plan is formed in the primary motor cortex, which is then executed by the release of efferent action potentials through the corticospinal tract that activate the relevant muscles. Studying this process, particularly the formation of the motor plan, provides valuable insight into the CNS. Additionally, we can compare the planning process with survivors of neurological injuries (*e.g.,* stroke) to study how the injuries influence motor planning.

We can examine motor planning by measuring changes in cortical activity during the interval between the cue to initiate movement and movement onset. Researchers will typically use transcranial electrical (Rossini et al., 1988) or magnetic (Bestmann & Duque, 2016; Chen et al., 1998; Ibanez et al., 2020) stimulation (TES or TMS, respectively) to stimulate a participant’s primary motor cortex just before a real (Copithorne et al., 2015; Davey et al., 1998; Leocani et al., 2000) or imaged (Facchini et al., 2002; Kumru et al., 2008; Zschorlich & Kohling, 2013) movement. Responses to the stimulation are recorded in the peripheral muscles groups that articulate the moving joint. These responses, called motor evoked potentials (MEPs), show the activity of the corticospinal tract (Bestmann & Duque, 2016). Previous studies using these approaches have shown that cortical excitability leading up to movement onset is characterized by initial inhibition, followed by an increase in excitability around 100 ms prior to movement onset (Chen & Hallett, 1999; Ibanez et al., 2020; Rossini et al., 1988). This initial inhibition is thought to allow the motor cortex to select between competing movement plans and prevent premature movements, while the subsequent increase in excitability corresponds to the facilitation of the selected motor plan and the inhibition of alternative plans (Bestmann & Duque, 2016; Labruna et al., 2014). Previous studies have also found that MEPs in muscles related to the movement are facilitated, while MEPs in unrelated or antagonist muscles are inhibited (Ganguly et al., 2021; Lebon et al., 2019; Reuter et al., 2015; Sohn & Hallett, 2004a, 2004b).

This facilitation/inhibition is thought to represent the plan formed by the motor cortex: facilitated muscles will be activated during the movement, while inhibited will not. Previous studies have also found that these processes are influenced by age (Reuter et al., 2015) and neurological disorders (Sohn & Hallett, 2004a).

However, these previous studies have only examined single degree of freedom movements (*e.g.,* wrist flexion/extension), and have not considered more complex, multi-joint movements. This distinction is particularly important when considering the upper extremity because it is a kinematically redundant system *i.e.,* different paths in the joint space can create the same path in the end-point space. This redundancy has several potential consequences on motor planning in the cortex. First, the distinction between muscles that are relevant and irrelevant to a movement are less clear. MEPs in muscles that seem irrelevant to the movement may be facilitated because the muscles serve a secondary function in the movement (*e.g.,* biceps brachii providing anti-gravity compensation during elbow extension). Second, different paths in the joint space require different muscle coordination, and therefore the representation of the motor plan in the cortex may vary depending on the movement. Finally, previous research has hypothesized that the CNS reduces the complexity of musculoskeletal control by synchronously activating groups of muscles in fixed patterns, called “muscle synergies” (Sherrington, 1910; Ting, 2007; Ting & McKay, 2007; Tresch & Jarc, 2009). If these synergies are indeed fixed, they may additionally influence corticospinal excitability during movement planning. Therefore, measuring changes in corticospinal excitability prior to the onset of multi-joint, upper extremity movements could reveal useful information regarding how the central nervous system plans and executes movement. The objective of this study was to examine how motor plans are expressed in the cortex prior to the onset of upper extremity movements. Specifically, we used single-pulse TMS to measure MEPs prior to the onset of a functional reaching task in several key upper extremity muscles. During the experiment, we varied muscle coordination by altering each participant’s initial arm posture so that different muscles were required to initiate the movement. We hypothesized that MEPs would be facilitated in all upper extremity muscles due to the multiple joints and muscles necessary to initiate the reach. Further, we hypothesized that corticospinal excitability would modulate with initial arm posture, as the arm posture would necessitate different muscle coordination to complete the reach.

## Methods and Materials

### Participants

20 adults (12 males, 8 females, age 22.1 yr ± 4.5 yr) with no history of major orthopedic or neurological conditions, injuries in their upper extremities, uncontrolled illnesses, and/or changes in the past 3 months participated in this study. Participants provided informed, written consent prior to participation, and all protocols received approval from the University of Michigan Institutional Review Board.

### Experimental Apparatus

Participants were seated at a table with a cup placed in front of them but just outside of their reach (Figure 1(A)). A computer monitor was also placed in front of the participant that displayed a green indicator light. Participants were instructed to wait for the green indicator to illuminate (*i.e.,* the “Go Cue”), and in response to the go cue, extend their dominant arm towards the target cup and extend their fingers, as if they were reaching out to grab it. Participants were instructed to react as quickly as possible to the go cue, but to move their arm towards the cup at a comfortable, self-selected pace. Participants were also instructed to keep their trunk stationary during the reach. The go cue was controlled by a custom program written in LabView (National Instruments Corp., Austin, TX USA) that could trigger a transcranial magnetic stimulator following an experimenter-defined time delay after the go cue (Figure 1(B)). When a participant’s reaction time was known (*i.e.*, the time between the go cue and movement onset), the delay could be used to deliver a stimulation at a desired time prior to movement onset.

**Figure 1.**
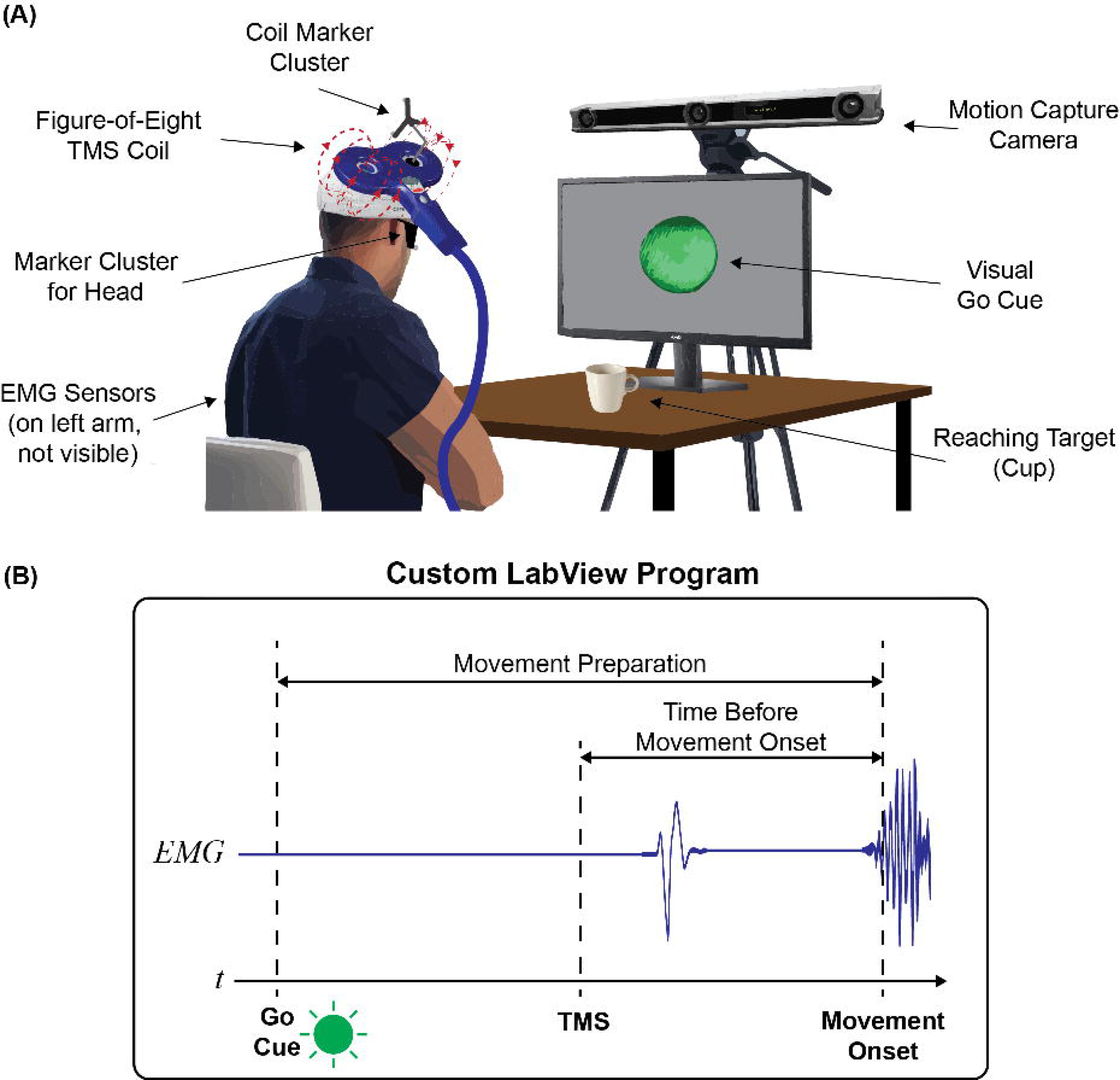
(A) The experimental apparatus. Each participant sat at a height-adjustable table facing a computer monitor. A cup was placed on the table in between the participant and the monitor, just outside of the participant’s reach. A green light was displayed on the monitor that would periodically light up. Participants were instructed to wait for the green light to illuminate, which was a “Go Cue” for the participants to extend their dominant arm (the left arm in the figure) forward as if they were going to grab the cup. During the experiment, a figure-of-eight transcranial magnetic stimulation (TMS) coil would periodically produce a single, monophasic magnetic pulse to stimulate the participant’s contralateral primary motor cortex. The responses to these stimulations, called MEPs, were measured using EMG sensors placed over key upper extremity muscles on the dominant arm. Marker clusters placed on the coil and on lenseless glasses worn by the participant were measured by a motion capture system to ensure consist coil placement. **(B)** TMS stimulation procedure: The go cue was controlled by a custom LabView program which triggered a TMS pulse following an experimenter-defined time delay. The time delay of the TMS pulse was configured so that it occurred at a desired time interval prior to movement onset.

Participants reached for the cup starting from one of two possible arm postures: “Elbow Down” and “Elbow Up” (Figure 2). In the “Elbow Down” position, the participants’ shoulder and forearm were in the neutral positions with the elbow flexed to 90 degrees. Their hand rested on their thigh with their fingers lightly flexed into a fist. In the “Elbow Up” position, the participants’ forearm and elbow rested on the table in front of them with their fingers lightly flexed into a fist in their sagittal plane. In this position, the shoulder was abducted, the elbow was flexed to 90 degrees, and the forearm was pronated. In the Elbow Up position, the height of the table was adjusted such that the participant’s elbow was just below the height of their shoulder.

**Figure 2.**
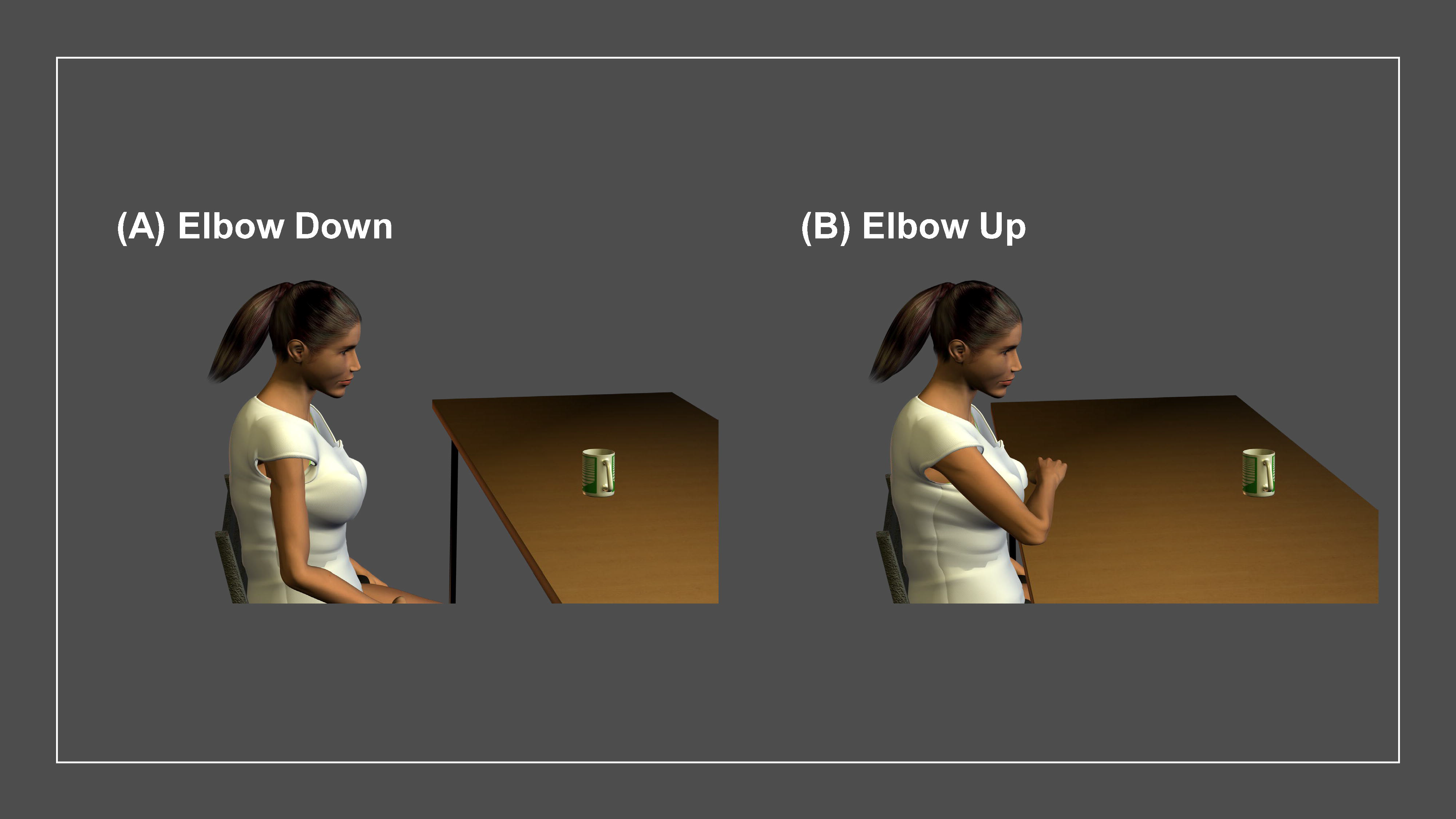
The two initial arm postures used in the experiment. **(A)** The Elbow Down posture. In this posture, the participant sat with their shoulder in the neutral position and elbow resting against their side. The elbow was flexed to 90°, the forearm was in the neutral position, and the hand rested on the participant’s ipsilateral thigh with the fingers lightly flexed into a fist. **(B)** The Elbow Up posture. In this posture, the participant sat with their shoulder abducted and their elbow resting on the table. The elbow was flexed to > 90° with the forearm pronated, the fingers flexed into light fist, and the hand in the sagittal plane.

We used these two postures because each required different muscles to initiate the reach. Specifically, the Elbow Down posture initially required shoulder flexion and elbow extension, with wrist and finger extension later in the reach to open the fingers. It is important to note, however, that gravity assisted elbow extension in this posture, therefore the biceps brachii provided anti-gravity assistance to control the elbow extension. Following this, we expected that the muscle coordination pattern immediately following movement onset would be dominated by the anterior deltoids, biceps brachii, and triceps brachii. The Elbow Up posture, on the other hand, initially required horizontal shoulder adduction and vertical shoulder abduction to rotate and lift the humerus, respectively, as well as elbow extension to extend the arm, and wrist extension to counteract gravity. Later in the reach, the participant would extend their wrist and fingers to grasp the cup. Thus, we expected that the muscle coordination just after movement onset in the Elbow Up posture would be dominated by the anterior and middle deltoids, triceps brachii, and wrist extensors. Therefore, if corticospinal excitability prior to movement onset reflects the subsequent muscle coordination, we expected that these muscle coordination patterns would emerge in the evoked responses prior to movement onset.

### Experimental Protocol

The experiment was split into two sections, one for each arm posture. The order of the postures was pseudorandomized for each participant. In the first section, participants performed a rest trial, a movement onset trial, and five reaching trials in the first arm posture. During the rest trial, we stimulated the participant’s contralateral motor cortex while the participant relaxed their arm. During the movement onset trial, participants reached for the cup as soon as they saw the go cue. During this trial, we measured the time between the go cue and the participant’s movement onset, and no stimulations were delivered. Following the movement onset trial, we performed five reaching trials. During the reaching trials, the participants reached for the cup in response to the go cue and received a stimulation prior to movement onset. In the first reaching trial, the stimulation time delay was set to 200 ms prior to movement onset. In the following reaching trials, we reduced the time between stimulation and movement onset in 50 ms increments until the last reaching trial, where the stimulation occurred 0 ms prior to movement onset. In the second section, participants repeated the rest and reaching trials in the second arm posture. Each trial consisted of ten stimulations/reaches. Following data collection, we recorded each participant’s maximum voluntary isometric contraction in shoulder flexion and abduction, elbow flexion and extension, and wrist flexion and extension while the participants’ arm were in the Elbow Down condition.

### Electromyography and TMS

During the experiment, we recorded each participant’s electromyography (EMG) to measure their responses to the TMS (*i.e.,* motor evoked potentials [MEPs]), as well as their muscle activity during reaches. To measure EMG, we placed single-use, Ag/AgCl electrodes (22 mm electrode spacing, NOROTRODE 20 Bipolar SEMG Electrodes, Myotronics, Kent, WA) over the muscle bellies of the anterior deltoid (AD), middle deltoid (MD), biceps brachii (BB), triceps brachii (TB), extensor digitorum superficialis (WE), and flexor digitorum superficialis (WF) of each participant’s dominant arm. Prior to electrode placement, the skin at each electrode site was cleaned with alcohol pads to minimize skin impedance, and electrode cream (Signacreme, Parker Laboratories, Inc., Fairfield, NJ) was applied to electrodes to increase conductivity between the skin and the electrode. Electrodes were fixed to the skin with the double-sided adhesive layer (applied to the electrode backing by the manufacturer) and further secured with tape. Electrodes were connected to preamplifiers (MA-422, Motion Lab Systems, Baton Rouge, LA) with snap connectors. Signals from each preamplifier were filtered with a 1000 Hz low-pass analog filter (MA300, Motion Lab Systems, Baton Rouge, LA) to eliminate signal aliasing, and then sampled at 2000 Hz with an 18-bit National Instruments Data Acquisition system (NI USB-6218, National Instruments Corp., Austin, TX USA). Sampled signals were recorded using a custom program written in LabView 2014 (National Instruments Corp., Austin, TX USA). Prior to data collection, the quality of each EMG signal was visually inspected to ensure proper electrode placement.

We performed TMS with a 70mm figure-of-eight coil connected to a monophasic magnetic stimulator (Magstim 200, Magstim, U.K.). The coil was positioned over the participant’s primary motor cortex contralateral to their dominant upper extremity. The coil was oriented to produce posterior-to-anterior current flow in cortex; the handle of the coil oriented ∼45° from the participant’s midline (pointed posterior). To find each participant’s hotspot (*i.e.,* the coil position and orientation producing the largest MEP), we initially placed the coil at the estimated hotspot (over the contralateral hemisphere, 3 cm lateral and anterior to the vertex), and made small adjustments. During hotspot finding, we looked for a location that elicited MEP in all muscles, however, if this was not possible, we used the location that produced the largest MEP in the middle deltoid. To ensure that we retained consistent coil position and orientation during data collection, we fixed a cluster of three retroreflective markers (9 mm diameter) to the coil and had participants wear custom, lenseless glasses with five retroreflective markers attached to them. The positions of both the coil and glasses marker clusters were tracked using an OptiTrack V120: TRIO camera and Motive Motion Capture Software (Version 1.8.0, 120Hz). The positions and orientations of the coil and the participant’s head, as well as the hotspot, were displayed to the experimenter during data collection using NeuRRoNav, a low-cost, open source software for navigated TMS (Rodseth et al., 2017). All stimulations were performed at 100% resting motor threshold (RMT), which was determined as the minimum threshold that produced MEPs more than 50% of the time and using the MTAT 2.0 program (MTAT 2.0)) (Awiszus & Borckardt, 2011). Threshold was determined by the threshold that produced MEPs in all muscles, but if this was not possible, we used the threshold that produced and MEP in the middle deltoid.

### Data Analysis

To validate that recorded muscle activity after movement onset reflected the hypothesized muscle coordination in the two postures, we examined the EMG signals from the first reaching trial using a custom program written in MATLAB (R2019b, MathWorks, Natick, MA). The raw EMG signals were first digitally filtered with a 20-500 Hz band-pass filter (zero-lag, 4^th^ order Butterworth) and then a 58-62 Hz band-rejection filter (zero-lag, 4^th^ order Butterworth) to eliminate surrounding electrical noise. We then deducted the mean from the filtered signals to remove the DC gain, rectified the signals, and smoothed them with a digital, 6 Hz low-pass filter (zero-lag, 4^th^ order Butterworth). The same filtering and smoothing process was applied to the recorded EMG during MVIC, and the smoothed MVIC was used to normalize the smoothed EMG from the reaching trial. Using the smoothed and normalized signals, we computed the ensemble average activation of the first 100 ms following movement onset, and then averaged over this 100 ms to get the average activity of each muscle just after movement onset for each participant. We averaged these values across participants to get the average muscle activity for each muscle in each arm posture.

To measure MEPs, each EMG signal was filtered using a zero-lag, second order bandpass Butterworth filter (passband: 0.25-1000 Hz). The filtered signals were separated into 150 ms segments (300 samples) following each stimulation, which we used to compute the ensemble average MEP for each muscle during each trial. The average MEP in a trial was defined as the peak-to-peak difference (*i.e.,* voltage maximum minus minimum) of this ensemble average.

Our primary outcome metric was change in MEP amplitude (ΔMEP) relative to resting at different stimulation timings (*e.g.,* 200 ms). To compute this change for a single participant, we computed the participant’s average MEP during a trial and deducted the average resting MEP recorded from the same arm posture. This difference was then normalized by the participant’s MVIC (the same value used for the muscle activation analysis). We averaged ΔMEP across all participants to get the average ΔMEP for each muscle during each reaching trial.

### Statistical Analysis

We performed our statistical analysis using IBM SPSS (Statistical Product and Service Solution, Version 27). To analyze our primary outcome metric, we performed a two-way repeated measures analysis of variance (ANOVA) for each muscle. The two factors for this analysis were stimulation timing (200, 150, 100, 50, and 0 ms), arm posture (Elbow Up and Elbow Down). A significant main or interaction effect was investigated by a post-hoc test with Sidak correction. We used 0.05 as the significance level for all analyses.

## Results

In general, the MEP of the proximal muscles (AD, MD, BB, and TB) were modulated by the timing of the stimulation (Figures 3). MEPs for these muscles remained near the resting MEP for stimulation timings 200, 150, and 100 ms prior to movement onset. At stimulation timings 50 and 0 ms prior to movement onset, MEPs in the proximal muscles increased above the resting MEP. The stimulation timing did not modulate the excitability of the distal muscles (WE and WF) as dramatically. These trends were observed in both arm postures. The BB showed the largest increases in MEP as the stimulation timing approached movement onset. The MD showed larger MEPs in the Elbow Up posture, while the BB showed larger MEPs in the Elbow Down posture. MEPs in the AD, TB, WE, and WF were similar between postures. These differences in MEP reflected the relative changes in muscle activity between postures just after movement onset, except for the TB, which showed larger activation in the Elbow Up posture (Figure 4).

**Figure 3.**
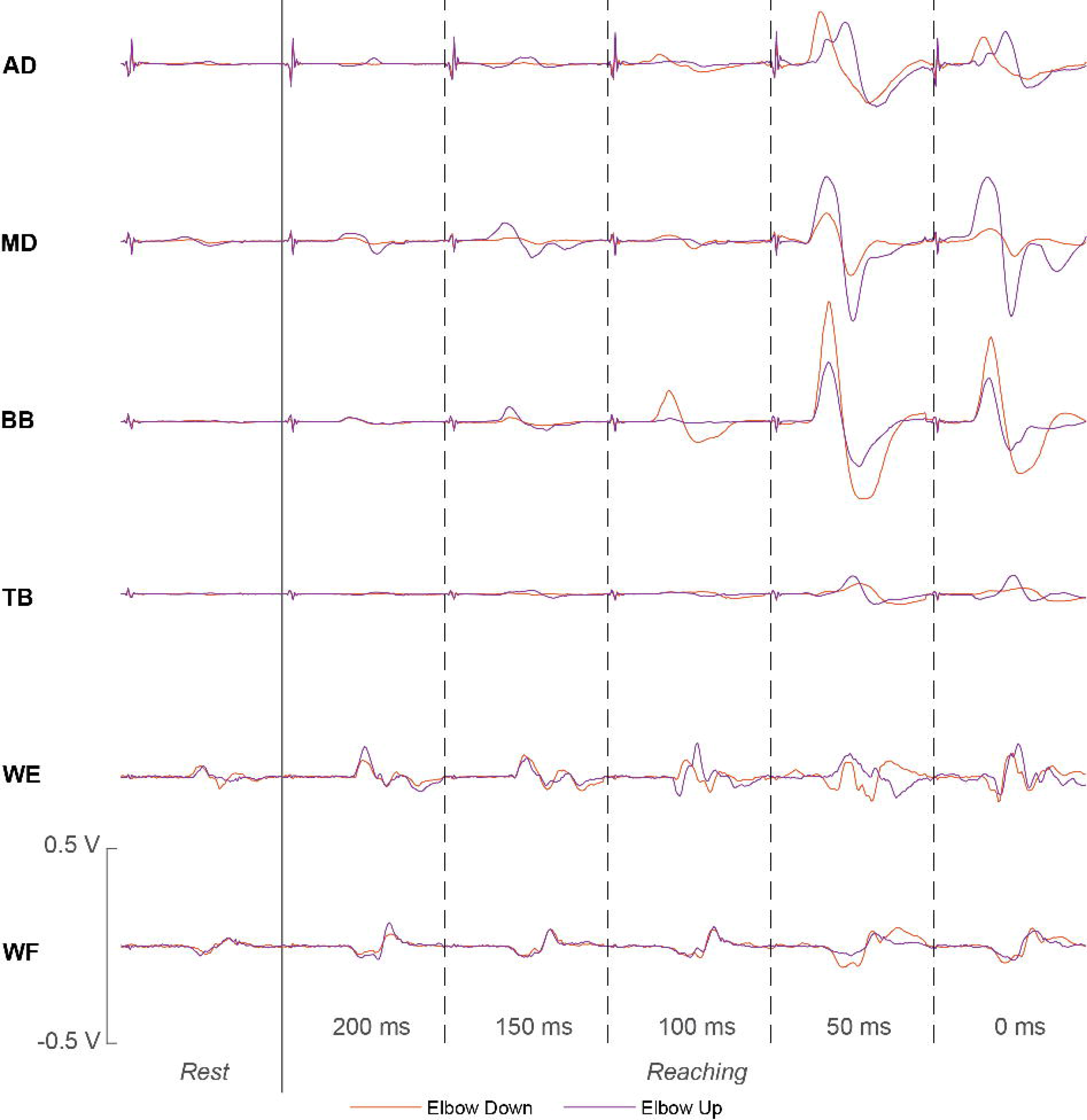
Ensemble average MEP from a typical participant in the measured upper extremity muscles and the in the two arm postures. Here, AD = Anterior Deltoid, MD = Middle Deltoid, BB = Biceps Brachii, TB = Triceps Brachii, WE = Wrist and Finger Extensors, and WF = Wrist and Finger Flexors.

**Figure 4.**
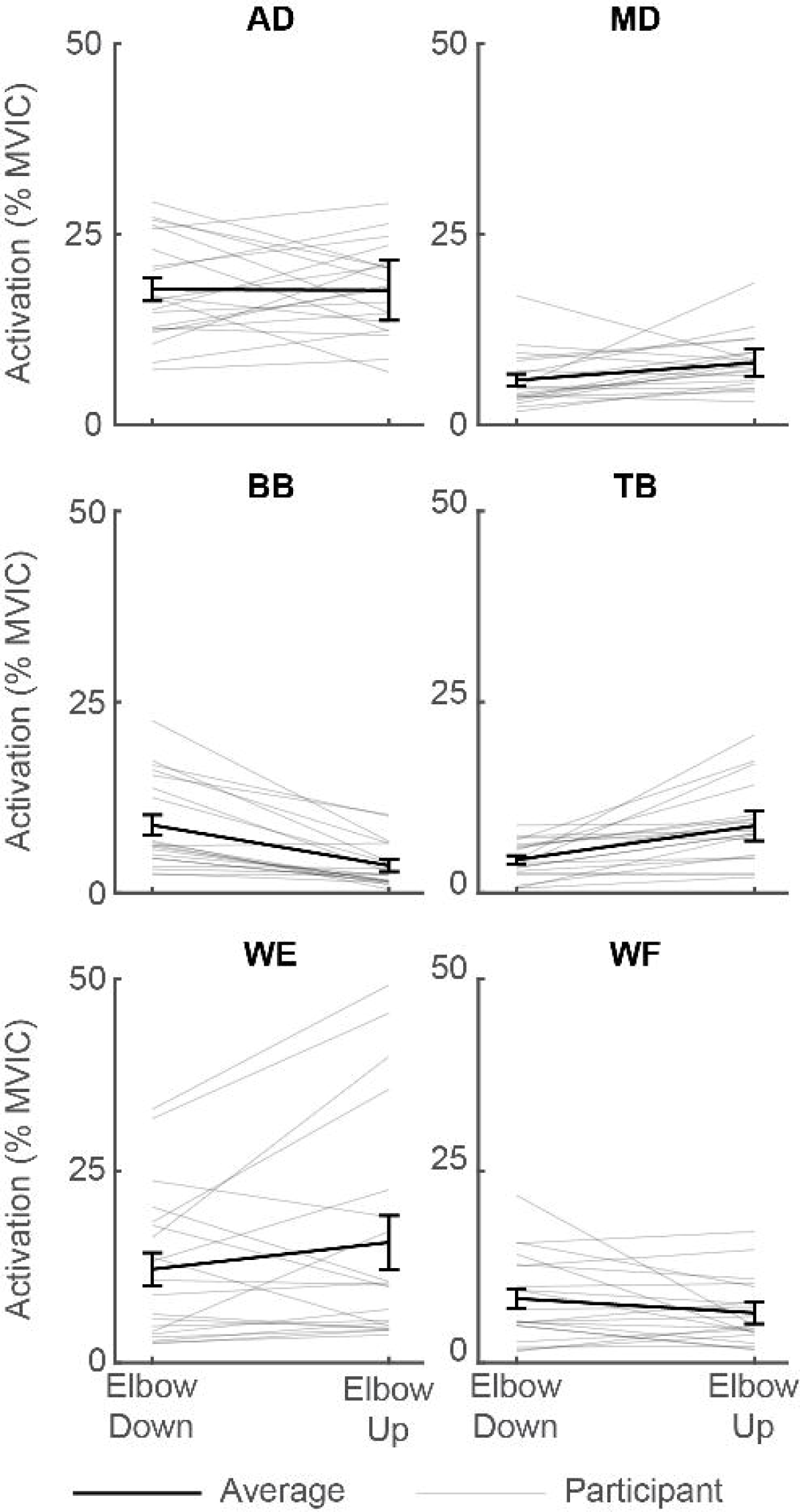
Participant and average muscle activation following movement onset between arm postures. Here, the error bars denote standard error of the mean.

We detected a significant main effect of timing in the ΔMEP of the anterior deltoid (AD, *p* = 0.004), middle deltoid (MD, *p <* 0.001), and triceps brachii (TB, *p* = 0.002) (Figure 5). Post-hoc analysis of the ΔMEP of the AD showed that ΔMEP at 200 ms differed from the ΔMEP at 100, 50, and 0 ms (*p* = 0.004, 0.005, and 0.003, respectively), ΔMEP at 150 ms differed from the ΔMEP at 50 and 0 ms (*p* = 0.009 and 0.004, respectively), ΔMEP at 100 ms differed from the ΔMEP at 50 and 0 ms (*p* = 0.009 and 0.004, respectively), and ΔMEP at 50 ms differed from the ΔMEP at 0 ms (*p* = 0.017). Post-hoc analysis of the ΔMEP of the MD showed that ΔMEP at 200 ms differed from the ΔMEP at 50 and 0 ms (*p* < 0.001 in both cases), ΔMEP at 150 ms differed from the ΔMEP at 50 and 0 ms (*p* < 0.001 in both cases), ΔMEP at 100 ms differed from the ΔMEP at 50 and 0 ms (*p* < 0.001 in both cases), and ΔMEP at 50 ms differed from the ΔMEP at 0 ms (*p* = 0.016). Post-hoc analysis of the ΔMEP of the TB showed that ΔMEP at 200 ms differed from the ΔMEP at 50 and 0 ms (*p* = 0.001 and < 0.001, respectively), ΔMEP at 150 ms differed from the ΔMEP at 50 and 0 ms (*p* < 0.001 in both cases), ΔMEP at 100 ms differed from the ΔMEP at 50 and 0 ms (*p* < 0.001 and = 0.001, respectively), and ΔMEP at 50 ms differed from the ΔMEP at 0 ms (*p* = 0.018).

**Figure 5.**
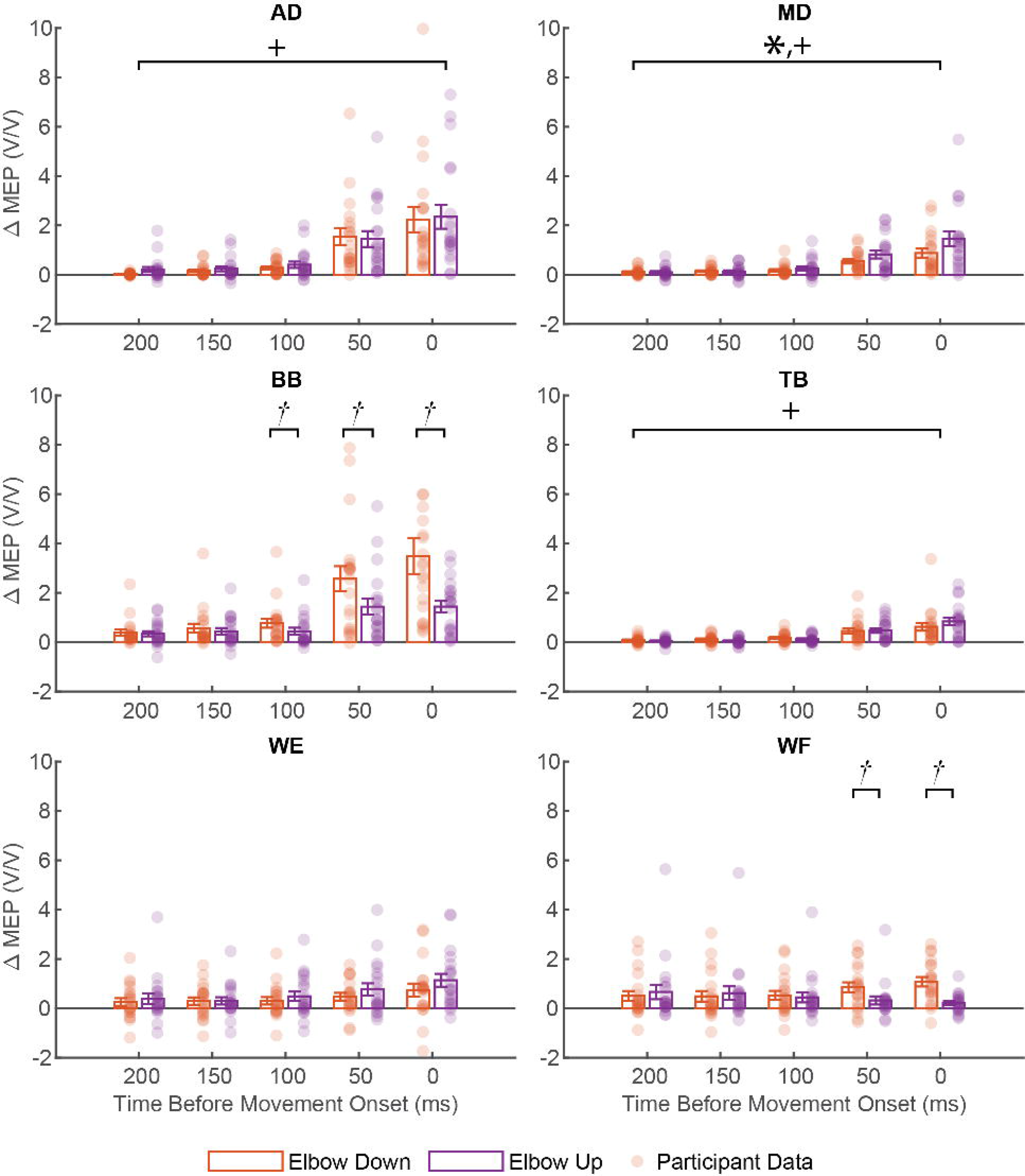
Participant and average ΔMEP in each arm posture and at each timing interval prior to movement onset. Here, the circle markers denote each participant’s trial ensemble average, the height of the bar denotes the trial average across all participants, and the error bars denote the standard error of the mean.

For the MD, the ΔMEP became larger in the Elbow Up posture than the Elbow Down as the stimulation timing approached movement onset. This trend, however, did not reach the level of significance (timing-by-posture interaction *p* = 0.157), although we did detect a significant main effect of posture (*p =* 0.038). The ΔMEP of the BB became larger in the Elbow Down posture than the Elbow Up as the stimulation timing approached movement onset. This trend did reach the level of significance (timing-by-posture interaction *p =* 0.013). Post-hoc analysis revealed that ΔMEP of the BB was significantly greater in the Elbow Down posture than the Elbow Up posture when the stimulation was delivered 100 ms, 50 ms, and 0 ms prior to movement onset (*p* = 0.012, < 0.001, and 0.005, respectively). In the Elbow Down posture, ΔMEP at 200 ms differed from the ΔMEP at 100, 50, and 0 ms (*p* = 0.016, 0.001, and 0.001, respectively), ΔMEP at 150 ms differed from the ΔMEP at 50, and 0 ms (*p* = 0.001 and 0.001, respectively), and ΔMEP at 100 ms differed from the ΔMEP at 50, and 0 ms (*p* < 0.001 and = 0.001, respectively). In the Elbow Up posture, ΔMEP at 200 ms differed from the ΔMEP at 50 and 0 ms (*p* = 0.005 and < 0.001, respectively), ΔMEP at 150 ms differed from the ΔMEP at 50, and 0 ms (*p* = 0.001 and 0.001, respectively), and ΔMEP at 100 ms differed from the ΔMEP at 50, and 0 ms (*p* = 0.001 and 0.003, respectively).

Regarding the distal muscles, we detected a significant timing-by-posture interaction (*p =* 0.014) in the ΔMEP of the WF. Specifically, the ΔMEP was significantly greater in the Elbow Down posture than the Elbow Up posture when the stimulation was delivered 50 ms and 0 ms prior to movement onsent (*p* = 0.023 and < 0.001, respectively). In the Elbow Down posture, the ΔMEP of the WF increased as the stimulation timing prior to movement onset moved towards 0 ms, while it decreased in the Elbow Up posture. This trend, however, was not significant, as the ΔMEP did not differ significantly across stimulation timings in either posture. We did not detect any significant main effects or interactions in WE.

## Discussion

The objective of this manuscript was to examine how motor planning is expressed in the cortex prior to upper extremity reaching tasks, and how this expression is influenced by kinematic redundancy. To examine this, we asked participants to perform a reaching task in response to a visual cue. Participants started from one of two arm postures that required different muscle coordination to complete the reach. Using TMS, we measured corticospinal excitability at several different upper extremity muscle and at several different time points prior to movement onset. We found that the excitability of all proximal muscles (AD, MD, BB, TB) increased as the stimulation timing approached movement onset. We also found that MEPs were largest in the MD and BB as the stimulation neared movement onset. Additionally, we found that this increase was modulated by arm posture. These results suggest that while changes corticospinal excitability are movement-specific, there is also a general excitability increase that is observable across many muscles, regardless of their contribution to the movement. These findings reveal valuable information regarding how the central nervous system plans movements.

We found that MEPs elicited close to movement onset (< 100 ms) increased in all proximal muscles, regardless of the initial posture of the upper extremity. This finding was somewhat surprising, as previous research has shown that MEPs during movement preparation are inhibited in irrelevant or antagonist muscles (Lebon et al., 2019; Reuter et al., 2015). It is likely that our findings differ from those of previous studies because the movements considered in this study include multiple degrees of freedom, while previous studies have only considered single-DOF movements. Following this, it is more difficult to discern what muscles should be considered irrelevant or antagonistic. For instance, we observed facilitation of biceps brachii MEPs, particularly in the Elbow Down posture, despite the task requiring elbow extension in both cases. However, in the Elbow Down posture, the biceps play a role in counteracting gravity to allow for a smoother, more controlled motion. Following this, it is possible that the general increase in excitability observed in all proximal muscles results from all proximal muscles serving a purpose in the subsequent movement, with some more obvious than others.

Interestingly, we found that MEPs measured in the distal upper extremity muscles were not facilitated and, in some cases, even inhibited (although not significantly). We believe that this occurred because of the time course of the reaching movement. Specifically, both reaching movements in this study are characterized by initial shoulder and elbow movement to extend the arm, followed by a delayed opening of the fingers to grasp the cup. As such, the proximal muscles are more important to the motor plan at movement onset, while the distal muscles are not recruited until later. However, it is currently unclear what this lack of facilitation/inhibition in the distal muscles represents at the level of motor planning in the cortex. For instance, it is possible that the lack of facilitation of MEP in the wrist muscles reflects a motor plan that, at the moment of measurement, is more focused on proximal muscles. Previous studies in serial motor tasks support this view, showing that when the cortex plans a sequence of tasks, all features of the sequence are developed in parallel prior to movement onset. The resulting movement corresponds to selective facilitation and inhibition of different features of the motor plan (Behmer & Crump, 2017; Behmer et al., 2023; Hurlstone et al., 2014). However, a competing theory is that preceding actions in a sequence act as cues for subsequent actions, triggering a new planning and action phase (Behmer et al., 2023; Shiffrin & Cook, 1978). As such, the observed response in the distal upper extremity muscles could then represent surround inhibition (Sohn & Hallett, 2004b) or subject’s inattention to their distal muscles at this stage in the movement. Future research is needed to determine the underlying neural mechanisms causing this observation.

Despite the observed MEP facilitation in all proximal muscles, we did find differences in MEPs between arm postures. These differences occurred in two proximal muscles (the biceps brachii and middle deltoids) and one distal muscle (wrist flexors, although not significantly). It is also important to note that these differences matched the differences in muscle coordination of the initiated movement between the two postures (Figure 4). Previous studies have shown that MEPs are facilitated during movement preparation in muscles relevant to the movement (Ganguly et al., 2021; Lebon et al., 2019; Reuter et al., 2015; Sohn & Hallett, 2004a, 2004b).

Our findings supplement these previous studies by showing that the magnitude of facilitation reflects the *importance* of the muscle in the subsequent movement (*e.g.,* a movement requiring more middle deltoid activation will show great middle deltoid MEP facilitation during planning). This is a finding unique to our study and arises from the inherent kinematic redundancy of the upper extremity. Because of the kinematic redundancy, it is possible to alter the contributions of muscles to complete the same functional task (*e.g.,* reaching for a cup) in multiple ways.

However, previous studies have only considered movements in one degree of freedom movements, which makes modulating the contribution of muscles much more difficult. Further testing with other postures and movements with other extremities is necessary to examine the full extent of how muscle coordination is reflected in the movement preparation phase.

As the stimulation timing approached movement onset, we found that excitability in both postures was characterized by large MEPs in the biceps brachii (BB) and smaller MEPs in the triceps brachii (TB). This result was surprising because reaching required elbow extension immediately following movement onset, and therefore one would expect higher excitability in TB (an elbow extensor) and lower excitability in BB (an elbow flexor). However, it is important to note that reaches in both postures required shoulder flexion and abduction. Therefore, the large excitability observed in the biceps brachii may be partially explained as an emergence of the “flexor synergy”. Conventionally, the flexor synergy is defined as the abnormal coupling of shoulder abduction and elbow flexion following a stroke (Dewald et al., 1995; Twitchell, 1951). Previous studies have shown that this coupling interferes with arm movements requiring elbow extension with shoulder flexion (Zackowski et al., 2004) or shoulder abduction (Sukal et al., 2006). Here, we observed a similar phenomenon: increased excitability of the BB during shoulder flexion and abduction, despite the necessity for concurrent elbow extension. This may suggest that the flexor synergy is present in humans without neurological injury, although to a lesser extent. These findings shed light onto the possible origin of abnormal joint coupling in stroke survivors, which is currently unclear. Researchers have shown that abnormal joint couplings following stroke may have a cortical origin (Gerachshenko et al., 2008; Krishnan & Dhaher, 2012; Yao et al., 2009), while others have shown that couplings may come from increased reliance on either ipsilateral cortical projections (Schwerin et al., 2008) and/or the reticulospinal tract (Kuypers, 1964; McPherson et al., 2018; McPherson & Dewald, 2022).

Evidence of the flexor synergy in the contralateral cortex in uninjured adults, however, supports the position that abnormal joint couplings after stroke has a cortical origin.

The findings in this study, when considered in conjunction with evidence that abnormal synergies in stroke survivors have a cortical origin, have interesting implications for use in clinical populations. Specifically, TMS could be used as a tool to examine the contribution of the flexor synergy to impairment or even a metric to judge the effectiveness of a therapy intervention in treating the flexor synergy. However, eliciting MEPs is notoriously difficult in stroke survivors. MEPs in stroke survivors can be produced more reliably using paired-pulse approaches (Schwerin et al., 2011), but many researchers still rely on the use of background contractions to increase MEP reliability (Krishnan & Dhaher, 2012). However, background contractions can confound examinations of abnormal muscle coordination because MEPs in the contracting muscle will be biased, and researchers must develop complex testing rigs to control for this bias (Krishnan & Dhaher, 2012; Schwerin et al., 2008). However, as this manuscript has shown, corticospinal excitability increases prior to the onset of multi-joint movements, and the excitability reflects the muscle coordination of the subsequent movement without being confounded by background contractions. Therefore, TMS prior to the onset of a multi-joint movement may be a suitable paradigm for facilitating MEPs in stroke survivors. However, it is unclear whether corticospinal excitability prior to movement onset is representative of corticospinal excitability during movement (Copithorne et al., 2015; Lockyer et al., 2021).

Therefore, future testing with members of the stroke population is necessary to fully establish this paradigm.

In summary, we used transcranial magnetic stimulation to elicit MEPs during preparation of a multi-joint, upper extremity movement. We examined two different initial upper extremity postures that would produce differing muscle coordination during the movement and found that the differences in coordination were present in the MEPs. We also found that MEPs were facilitated in all proximal muscles as stimulations were delivered closer to movement onset.

Additionally, we found that MEPs expression reflected the flexor synergy, suggesting that the flexor synergy typically observed in stroke survivors may have a cortical origin. These results reveal interesting information regarding the mechanisms by which the motor cortex plans movements and may have utility for researchers examining muscle coordination in stroke survivors.

## Acknowledgements

This work was partly supported by grants from the National Science Foundation (Grant #’s 1804053 and 1256260), the National Institutes of Health (Grant # R01-EB019834), and the NICHD National Center of Neuromodulation for Rehabilitation (Grant # P2C-HD086844)

## Conflict of Interests Statement

None of the authors have any conflicts of interest.

## Notes

### Competing Interest Statement

The authors have declared no competing interest.

## References

1. Awiszus, F., & Borckardt, J. (2011). TMS motor threshold assessment Tool (MTAT 2.0). In ClinicalResearcher.org

2. Behmer, L. P., & Crump, M. J. (2017). The dynamic range of response set activation during action sequencing. J Exp Psychol Hum Percept Perform, 43(3), 537–554. https://doi.org/10.1037/xhp0000335

3. Behmer, L. P., Crump, M. J. C., & Jantzen, K. J. (2023). Motor-Evoked Potentials for Early Individual Elements of an Action Sequence During Planning Reflect Parallel Activation Processes. Motor Control, 1-20. https://doi.org/10.1123/mc.2022-0106

4. Bestmann, S., & Duque, J. (2016). Transcranial Magnetic Stimulation: Decomposing the Processes Underlying Action Preparation. Neuroscientist, 22(4), 392–405. https://doi.org/10.1177/1073858415592594

5. Chen, R., & Hallett, M. (1999). The time course of changes in motor cortex excitability associated with voluntary movement. Can J Neurol Sci, 26(3), 163–169. https://doi.org/10.1017/s0317167100000196

6. Chen, R., Yaseen, Z., Cohen, L. G., & Hallett, M. (1998). Time course of corticospinal excitability in reaction time and self-paced movements. Ann Neurol, 44(3), 317–325. https://doi.org/10.1002/ana.410440306

7. Copithorne, D. B., Forman, D. A., & Power, K. E. (2015). Premovement Changes in Corticospinal Excitability of the Biceps Brachii are Not Different Between Arm Cycling and an Intensity-Matched Tonic Contraction. Motor Control, 19(3), 223–241. https://doi.org/10.1123/mc.2014-0022

8. Davey, N. J., Rawlinson, S. R., Maskill, D. W., & Ellaway, P. H. (1998). Facilitation of a hand muscle response to stimulation of the motor cortex preceding a simple reaction task. Motor Control, 2(3), 241–250. https://doi.org/10.1123/mcj.2.3.241

9. Dewald, J. P., Pope, P. S., Given, J. D., Buchanan, T. S., & Rymer, W. Z. (1995). Abnormal muscle coactivation patterns during isometric torque generation at the elbow and shoulder in hemiparetic subjects. Brain, 118 ( *Pt 2*), 495–510. https://doi.org/10.1093/brain/118.2.495

10. Facchini, S., Muellbacher, W., Battaglia, F., Boroojerdi, B., & Hallett, M. (2002). Focal enhancement of motor cortex excitability during motor imagery: a transcranial magnetic stimulation study. Acta Neurol Scand, 105(3), 146–151. https://doi.org/10.1034/j.1600-0404.2002.1o004.x

11. Ganguly, J., Kulshreshtha, D., Almotiri, M., & Jog, M. (2021). Muscle Tone Physiology and Abnormalities. Toxins (Basel), 13(4). https://doi.org/10.3390/toxins13040282

12. Gerachshenko, T., Rymer, W. Z., & Stinear, J. W. (2008). Abnormal corticomotor excitability assessed in biceps brachii preceding pronator contraction post-stroke. Clin Neurophysiol, 119(3), 683–692. https://doi.org/10.1016/j.clinph.2007.11.004

13. Hurlstone, M. J., Hitch, G. J., & Baddeley, A. D. (2014). Memory for serial order across domains: An overview of the literature and directions for future research. Psychol Bull, 140(2), 339–373. https://doi.org/10.1037/a0034221

14. Ibanez, J., Hannah, R., Rocchi, L., & Rothwell, J. C. (2020). Premovement Suppression of Corticospinal Excitability may be a Necessary Part of Movement Preparation. Cereb Cortex, 30(5), 2910–2923. https://doi.org/10.1093/cercor/bhz283

15. Krishnan, C., & Dhaher, Y. (2012). Corticospinal responses of quadriceps are abnormally coupled with hip adductors in chronic stroke survivors. Exp Neurol, 233(1), 400–407. https://doi.org/10.1016/j.expneurol.2011.11.007

16. Kumru, H., Soto, O., Casanova, J., & Valls-Sole, J. (2008). Motor cortex excitability changes during imagery of simple reaction time. Exp Brain Res, 189(3), 373–378. https://doi.org/10.1007/s00221-008-1433-6

17. Kuypers, H. G. (1964). The Descending Pathways to the Spinal Cord, Their Anatomy and Function. Prog Brain Res, 11, 178–202. https://doi.org/10.1016/s0079-6123(08)64048-0

18. Labruna, L., Lebon, F., Duque, J., Klein, P. A., Cazares, C., & Ivry, R. B. (2014). Generic inhibition of the selected movement and constrained inhibition of nonselected movements during response preparation. J Cogn Neurosci, 26(2), 269–278. https://doi.org/10.1162/jocn_a_00492

19. Lebon, F., Ruffino, C., Greenhouse, I., Labruna, L., Ivry, R. B., & Papaxanthis, C. (2019). The Neural Specificity of Movement Preparation During Actual and Imagined Movements. Cereb Cortex, 29(2), 689–700. https://doi.org/10.1093/cercor/bhx350

20. Leocani, L., Cohen, L. G., Wassermann, E. M., Ikoma, K., & Hallett, M. (2000). Human corticospinal excitability evaluated with transcranial magnetic stimulation during different reaction time paradigms. Brain, 123 ( *Pt 6*), 1161–1173. https://doi.org/10.1093/brain/123.6.1161

21. Lockyer, E. J., Compton, C. T., Forman, D. A., Pearcey, G. E., Button, D. C., & Power, K. E. (2021). Moving forward: methodological considerations for assessing corticospinal excitability during rhythmic motor output in humans. J Neurophysiol, 126(1), 181–194. https://doi.org/10.1152/jn.00027.2021

22. McPherson, J. G., Chen, A., Ellis, M. D., Yao, J., Heckman, C. J., & Dewald, J. P. A. (2018). Progressive recruitment of contralesional cortico-reticulospinal pathways drives motor impairment post stroke. J Physiol, 596(7), 1211–1225. https://doi.org/10.1113/JP274968

23. McPherson, L. M., & Dewald, J. P. A. (2022). Abnormal synergies and associated reactions post-hemiparetic stroke reflect muscle activation patterns of brainstem motor pathways. Front Neurol, 13, 934670. https://doi.org/10.3389/fneur.2022.934670

24. Reuter, E. M., Behrens, M., & Zschorlich, V. R. (2015). Age-related differences in corticomotor facilitation indicate dedifferentiation in motor planning. Exp Gerontol, 65, 79–84. https://doi.org/10.1016/j.exger.2015.03.008

25. Rodseth, J., Washabaugh, E. P., & Krishnan, C. (2017). A novel low-cost approach for navigated transcranial magnetic stimulation. Restor Neurol Neurosci, 35(6), 601–609. https://doi.org/10.3233/RNN-170751

26. Rossini, P. M., Zarola, F., Stalberg, E., & Caramia, M. (1988). Pre-movement facilitation of motor-evoked potentials in man during transcranial stimulation of the central motor pathways. Brain Res, 458(1), 20–30. https://doi.org/10.1016/0006-8993(88)90491-x

27. Schwerin, S., Dewald, J. P., Haztl, M., Jovanovich, S., Nickeas, M., & MacKinnon, C. (2008). Ipsilateral versus contralateral cortical motor projections to a shoulder adductor in chronic hemiparetic stroke: implications for the expression of arm synergies. Exp Brain Res, 185(3), 509–519. https://doi.org/10.1007/s00221-007-1169-8

28. Schwerin, S. C., Yao, J., & Dewald, J. P. (2011). Using paired pulse TMS to facilitate contralateral and ipsilateral MEPs in upper extremity muscles of chronic hemiparetic stroke patients. J Neurosci Methods, 195(2), 151–160. https://doi.org/10.1016/j.jneumeth.2010.11.021

29. Sherrington, C. S. (1910). Flexion-reflex of the limb, crossed extension-reflex, and reflex stepping and standing. J Physiol, 40(1-2), 28–121. https://doi.org/10.1113/jphysiol.1910.sp001362

30. Shiffrin, R. M., & Cook, J. R. (1978). Short-term forgetting of item and order information. Journal of Verbal Learning and Verbal Behavior, 17(2), 189–218.

31. Sohn, Y. H., & Hallett, M. (2004a). Disturbed surround inhibition in focal hand dystonia. Ann Neurol, 56(4), 595–599. https://doi.org/10.1002/ana.20270

32. Sohn, Y. H., & Hallett, M. (2004b). Surround inhibition in human motor system. Exp Brain Res, 158(4), 397–404. https://doi.org/10.1007/s00221-004-1909-y

33. Sukal, T. M., Ellis, M. D., & Dewald, J. P. (2006). Source of work area reduction following hemiparetic stroke and preliminary intervention using the ACT3D system. Conf Proc IEEE Eng Med Biol Soc, 2006, 177–180. https://doi.org/10.1109/IEMBS.2006.259311

34. Ting, L. H. (2007). Dimensional reduction in sensorimotor systems: a framework for understanding muscle coordination of posture. Prog Brain Res, 165, 299–321. https://doi.org/10.1016/S0079-6123(06)65019-X

35. Ting, L. H., & McKay, J. L. (2007). Neuromechanics of muscle synergies for posture and movement. Curr Opin Neurobiol, 17(6), 622–628. https://doi.org/10.1016/j.conb.2008.01.002

36. Tresch, M. C., & Jarc, A. (2009). The case for and against muscle synergies. Curr Opin Neurobiol, 19(6), 601–607. https://doi.org/10.1016/j.conb.2009.09.002

37. Twitchell, T. E. (1951). The restoration of motor function following hemiplegia in man. Brain, 74(4), 443–480. https://doi.org/10.1093/brain/74.4.443

38. Yao, J., Chen, A., Carmona, C., & Dewald, J. P. (2009). Cortical overlap of joint representations contributes to the loss of independent joint control following stroke. Neuroimage, 45(2), 490–499. https://doi.org/10.1016/j.neuroimage.2008.12.002

39. Zackowski, K. M., Dromerick, A. W., Sahrmann, S. A., Thach, W. T., & Bastian, A. J. (2004). How do strength, sensation, spasticity and joint individuation relate to the reaching deficits of people with chronic hemiparesis? Brain, 127(Pt 5), 1035–1046. https://doi.org/10.1093/brain/awh116

40. Zschorlich, V. R., & Kohling, R. (2013). How thoughts give rise to action - conscious motor intention increases the excitability of target-specific motor circuits. PLoS One, 8(12), e83845. https://doi.org/10.1371/journal.pone.0083845

